# SELECTION OF ANTI-NUCLEAR ANTIGEN (ANA) REACTIVE B CELLS IN SYSTEMIC LUPUS ERYTHEMATOSUS

**DOI:** 10.1101/2025.01.08.631945

**Authors:** Yemil Atisha-Fregoso, Wenzhao Meng, Aaron M. Rosenfeld, Fang Liu, Thomas MacCarthy, Scott Feltman, Cynthia Aranow, Meggan Mackay, Chloe Terestchenko, Moriah Dunn, Matthew Scharff, Eline T. Luning Prak, Betty Diamond

## Abstract

**Objective:** Autoreactive B cells that recognize nuclear antigens are normally present in healthy individuals and patients with systemic lupus erythematosus (SLE), yet their activation and the production of IgG autoantibodies is a hallmark of SLE. The selection process and regulation of these cells in patients with SLE has not been completely understood. To gain insights into tolerance checkpoints and the developmental trajectories of autoreactive clones, we studied the BCR sequences from thousands of anti-nuclear antigen binding (ANA)+ and ANA-B cells from patients with SLE.

**Methods:** From a cohort of 13 patients with SLE, we identified and isolated ANA+ and ANA-B cells by flow cytometry using a method based on their binding to nuclear extracts. We sequenced B cell receptor (BCR) heavy chain variable regions and investigated the features of the IgH repertoire of ANA+ and ANA-B cells from naïve, memory and age-associated B cells (ABCs), and from total plasmablasts.

**Results:** The frequency of ANA+ B cells was similar in ABCs and naive B cells and higher in both than in memory B cells. We observed preferential usage of some VH (IGHV1-18, IGHV3-21, IGHV3-23|3-23D, IGHV4-34, IGHV4-39 and IGHV4-59) and VJ genes (IGHJ4 and IGHJ6) in B cells from these patients. ANA+ naïve and ANA+ ABCs used different gene segments and have longer CDR3 sequences than ANA+ memory B cells and ANA-subsets. ANA+ ABCs and memory B cells have a lower frequency of somatic hypermutation (SHM) and less activation induced deaminase (AID) targeting to WRC hotspots compared with their ANA-counterparts. Patients with active disease have a lower frequency of SHM in ANA+ ABCs and memory B cells and in ANA-ABCs.

**Conclusion:** Compared to memory B cells, ABCs are enriched in autoreactivity. Our results suggest that there is an immune checkpoint that restricts the differentiation of ANA+ naïve B cells into memory B cells and that ANA+ ABCs originate from ANA+ naïve B cells. Lower frequencies of SHM in antigen experienced ANA+ B cells, and particularly ANA+ ABCs, suggest that these cells might be generated through an extrafollicular (EF) pathway, and that in patients with active SLE there is more EF activation.

## INTRODUCTION

Antibodies expressed on the B cell membrane are termed B cell receptors (BCR). They are composed of two identical heavy chains and two identical light chains and allow the B cell to respond to antigen in the immediate environment. BCR diversity, critical for enabling a B cell response to a universe of antigens, is generated by recombination of variable (V), diversity (D) and joining (J) gene segments, junctional modifications and different combinations of antibody heavy and light chain pairs. The third complementarity determining region (CDR3) in the heavy chain, which spans the junctions between the V and J segments, is the most diverse region of the antibody. In addition to V(D)J recombination, further diversification occurs in mature B cells by somatic hypermutation (SHM) of the V gene. Due to the stochastic nature of these diversification processes, autoreactive receptors are frequently generated. Some B cells with autoreactive BCRs are subject to central mechanisms of tolerance, such as deletion or receptor editing [1,2]; nevertheless, in healthy subjects without any evidence of autoimmune disease, as many as 10 to 20% of circulating mature B cells are autoreactive [3–5]. Peripheral mechanisms of tolerance prevent the activation of autoreactive naïve B cells into pathogenic memory B cells and/or antibody forming cells (including plasma cells). Some or all of these tolerance checkpoints are compromised in individuals with autoimmune diseases [6,7].

Systemic lupus erythematosus (SLE) is an autoimmune disease that is characterized by the production of multiple pathogenic antinuclear autoantibodies (ANA) [8]. While the autoreactive B cells that produce these autoantibodies are considered central to SLE pathogenesis, their developmental origins remain unclear. Some studies indicate that patients with SLE have alterations in selection of autoreactive B cells during the development of the mature immunocompetent B cell repertoire prior to the formation of immune responses [9]. Other studies suggest that tolerance is broken during B cell activation due to B cell intrinsic and extrinsic differences in activation thresholds [6]. SHM plays a crucial role in the acquisition of autoreactivity in antigen-activated B cells as they proliferate and differentiate in the germinal center (GC) [10–12] . Other studies point to altered regulation of B cells as they differentiate to plasma cells, through extrafollicular (EF) immune activation [13–15].

Previous studies evaluated the BCR repertoire of blood B cells in patients with SLE without regard to their antigenic specificity [16–20]. Others have investigated the self-reactive status and BCR sequence of single B cell clones in patients with SLE; such studies are labor intensive and limited by the number of analyzed B cells [9,5]. No study so far has investigated the characteristics of the repertoire of large numbers of ANA+ and ANA- B cells in patients with SLE; the closest approximation is a study of cells bearing the 9G4 idiotype, which are presumed to be autoreactive [21,22].

We developed a method to distinguish ANA+ and ANA-B cells by flow cytometry based on their binding to nuclear extract [4]. Using this method, we explored the BCR heavy chain variable region sequence, and characterized features of the IgH repertoire of thousands of ANA+ and ANA-B cells. In addition to stratifying the circulating SLE repertoire by ANA reactivity, our analysis focuses on different B cell subsets: naïve, memory, age-associated B cells (ABCs), and plasmablasts. Plasmablasts are activated B cells that undergo cell division, secrete antibodies, and can migrate to the bone marrow where they differentiate further into plasma cells [23]. ABCs are memory-like cells that accumulate with age, exhibit sexual dimorphism in lupus mice [24] and respond to innate signals, such as TLR7 and TLR9 [25], which are relevant to the recognition of ANA [26]. There is evidence that supports an EF origin of ABCs [27,28]. Plasmablasts and ABCs are of particular interest in SLE, as circulating levels of these cells tend to increase during disease flares [29,30]. We focused on these subsets to gain insights into tolerance checkpoints and the developmental trajectories of autoreactive clones in human SLE.

## METHODS

### Patients

The SLE cohort analyzed in this cross-sectional study was recruited from the Northwell Health Lupus Center of Excellence. All patients were ≥ 18 years old, and fulfilled either the American College of Rheumatology or Systemic Lupus International Collaborating Clinics criteria for SLE [31,32]. Clinical and laboratory data were collected at the same time as the blood draw. Clinical activity was evaluated using the Systemic Lupus Disease Activity Index (SLEDAI)-2000 [33], clinical SLE Disease Activity Index (cSLEDAI) (which does not take serology into account) [34] and Physician Global Assessment (PGA) [35] scores. All subjects provided written informed consent. The study was approved by the Northwell Health Institutional Review Board. Overall study design is shown in Figure 1A.

**FIGURE 1.**
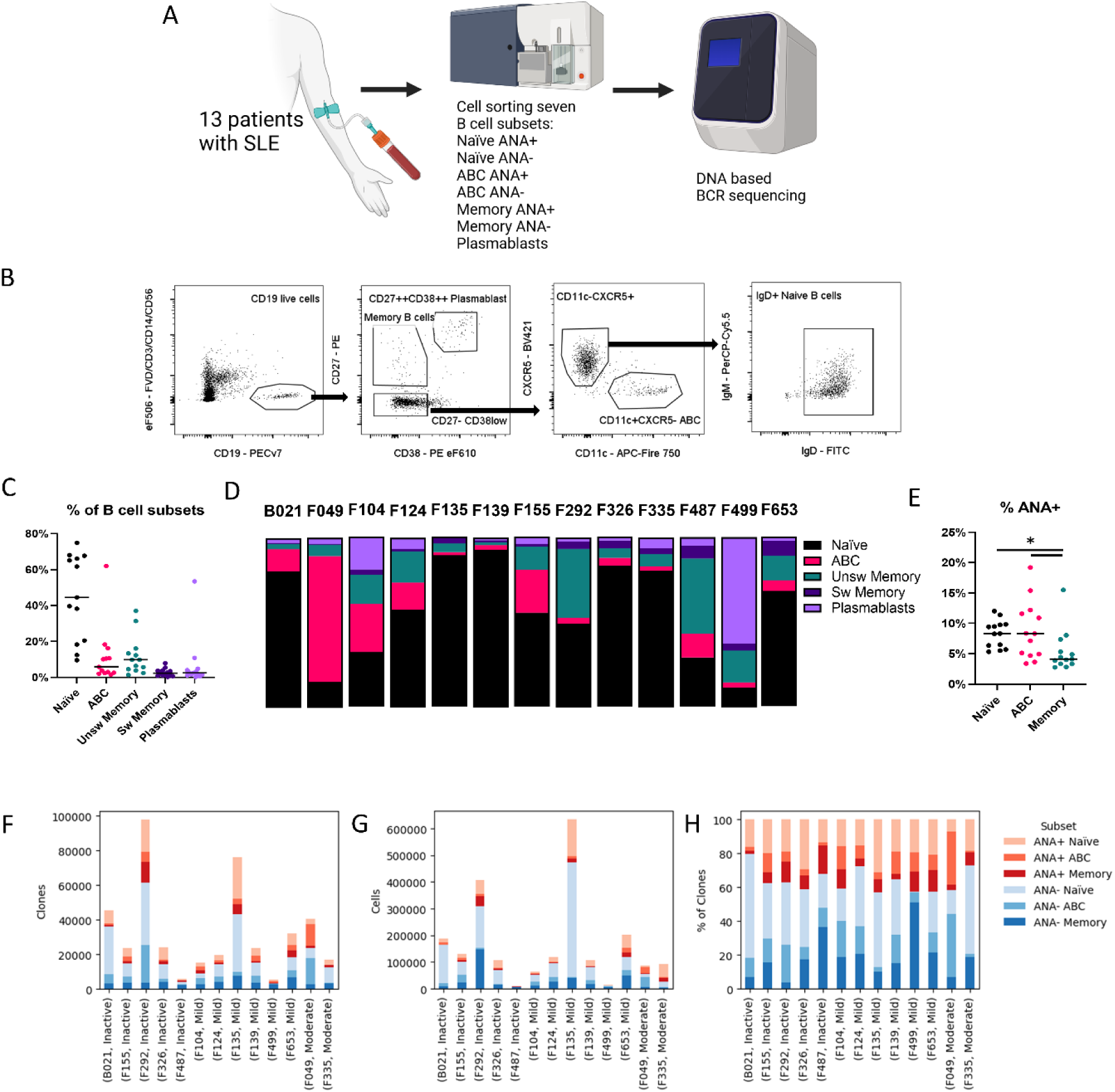
Study design and characteristics of B cells in patients with SLE. A) Overview of the study design. Peripheral blood samples from 13 patients with SLE were obtained, stained with surface antibodies and nuclear extract as described in methods, and seven subsets were isolated by FACS sorting. Cells were immediately lysed and DNA-based BCR sequencing was performed for further analysis. The sorter and sequencer depicted are for illustrative purposes only and do not reflect the specific equipment used in these experiments. B) Gating strategy for the identification of four main populations: naïve, memory, ABCs and plasmablasts. For the first three populations ANA+ and ANA-were identified and isolated. C) Proportion of B cell subsets identified by flow cytometry in the cohort. Each dot represents an individual patient. D) Relative distribution of B cell subsets in each individual patient. Each bar represents 100% of B cells for that patient. E) Proportion of ANA+ B cells in naïve, ABCs and memory B cells. Each dot represents an individual patient. F) Absolute number of clones obtained from each subset per individual donor after BCR sequencing. G) Absolute number of sorted cells from each subset per individual donor. H) Proportion of clones that belong to each subset per patient. Disease activity was defined according to PGA as described in methods. Statistics. Pairwise comparisons were performed between each pair of subsets using Wilcoxon signed rank test, and the false discovery rate approach was used to control for multiple comparisons, as described in methods. * adjusted p < 0.05. ANA: Antinuclear antigens. BCR: B cell receptor. PGA: Physician Global Assessment.

### B cell processing and ANA staining

Thirty to forty mL of blood were obtained from each patient by peripheral venipuncture. Peripheral blood mononuclear cells (PBMCs) were isolated by gradient density separation with Ficoll-Paque (GE healthcare) less than 4 hours after blood draw. Nuclear extracts from HeLa cells were obtained as previously described [4]. Briefly, nuclei from HeLa cells were isolated with the Nuclei EZ Lysis Kit (Sigma, St Louis, Mo). Nuclei were fragmented by vortexing with 0.5 mm glass beads (Scientific Industries, Bohemia, NY), and biotinylated using EZLink-Sulfo-NHS-LC-biotin (Thermo Scientific, Waltham, Ma). Cells were incubated with nuclear extract diluted in 1.5% nonfat dry milk (LabScientific, Highlands NJ) in Hank’s balanced salt solution. After thorough washing, cells were stained in HBSS plus 2% FBS containing 1 ug/mL streptavidin-allophycocyanin (Biolegend, San Diego, Ca) and fluorochrome labeled monoclonal antibodies (full list of antibodies is provided in supplementary methods). Dead cells were excluded with eFluor 506–labeled fixable viability dye (eBioscience, San Diego, Ca). After fixation and permeabilization procedure required for ANA staining of plasmablasts, the cellular DNA is of low-quality making identification of the IgG repertoire difficult; we therefore isolated total plasmablasts for BCR analysis.

### Immunophenotyping

B cell subset gating is shown in Figure 1B. B cells were defined as CD19+ and viability dye negative. We identified and sorted four B cell subsets: naïve (CD27-, CD38int, IgD+), memory (CD27+CD38int), ABC (CD11c+CXCR5-), and circulating plasmablasts (CD27++CD38++). Due to the limited number of subsets that we were able to sort simultaneously and the low number of cells in each sample, we combined unswitched (IgD+) and switched (IgD-) memory B cells (calling the combined population memory B cells). For cell sorting, the sample was divided in two: the first aliquot (two thirds of the volume) was used to sort ANA+ subsets and plasmablasts, while the second aliquot (one third of the volume) was used to sort ANA-subsets and also plasmablasts (which were not separated based on ANA binding).

### IgH Sequencing

Cells were sorted directly into lysis buffer (Qiagen cat. # 158113) and shipped to the University of Pennsylvania, where they were subjected to IgH bulk sequencing from genomic DNA using primers in FR1 and JH, as described previously [36,37]. Two biological replicates were generated and sequenced per sample except for samples with very low cell inputs (denoted with asterisks in Supplemental Table 4). Sequencing was performed on an Illumina MiSeq instrument in the Human Immunology Core facility at the University of Pennsylvania using Illumina 2×300 bp paired end kits (Illumina cat. # MS-102-3003).

### AIRR-seq data processing and additional filtering

Raw reads from the Illumina MiSeq were quality controlled, annotated, and grouped into clones as described previously [38]. Further clone collapsing was performed to group identical CDR3 amino acid sequences regardless of VH/JH, to capture additional clonal sequence variants. However, this CDR3 nucleotide collapsing method would occasionally introduce clonally unrelated sequences with very different VH genes. Therefore, sequence variants that contained 25 or more mutations compared to the preceding vertical node were removed. To minimize the effects of barcode hopping and cross-sample contamination, clones that were shared between different individuals at a ratio of 10:1 or higher were removed from the individual with the lower copy fraction. For clonal lineage and clonal overlap analysis (Venn Diagrams), clones with a subject-level copy number of 5 or fewer sequences were removed from the analysis. Clonal lineages were further filtered based on copy number of individual sequence variants (a minimum of 2 copies per node was required) to remove probable sequencing errors. Ete3 [39] was used to plot individual lineages. Only clones with productive rearrangements were analyzed in this paper.

### Mutation Targeting Analysis

Mutation analysis was performed as previously described [40,41]. All analyses were performed using custom R scripts. IGHV germline genes were obtained from IMGT [42]. The matchPattern function, part of the Biostrings library, was used to identify hotspot positions. We used the dominant, consensus sequence found for each clone and used this as the reference for all variant mutation calls. To reduce noise cause by sequencing artifacts, we only included clones with more than five members and limited the analysis of sequences with 3 to 16 mutations. After identification of hotspot profiles for each IGHV germline gene, to evaluate whether mutations in the sequences were targeted to hotspots or not, we assumed that the mutations occurred in the sequence background. To evaluate mutational enrichment to activation induced deaminase (AID) hotspots, we added up the mutations from each unique sequence, then considered the context in which they occurred (hotspot or non-hotspot). We compared the ratio of mutations targeting hotspots (AID targeting ratio) according to the formula: Observed = Number of mutations at WRC/ Total mutations. Expected = Number of WRC sites / sequence length. Then we calculated observed / expected for each clone, and calculated the average of the ratio for all clones for each V gene separately for ANA-vs ANA+.

### Statistical Analyses

Individual variables representing a point estimate of the value of a factor for each patient were compared using two-tailed Wilcoxon signed-rank test. To control the false discovery rate at 0.05 and to define significance, we used the Benjamini-Hochberg method [43]. When P values are mentioned, they represent adjusted P value.

Principal Component Analysis (PCA) of V Gene Usage. To analyze the patterns of V gene usage of the most prevalent genes across different B cell subsets, we performed Principal Component Analysis (PCA). The data were scaled and centered prior to PCA computation. PCA was performed using the prcomp function in R.

Analyses of CDR3 Length Distribution. To analyze the distribution of CDR3 lengths across different B cell populations, and to adjust for the inherent clustering caused by the donor, we employed a Generalized Linear Mixed Model (GLMM) approach. We considered B cell subpopulation a fixed effect and the individual a random effect. We pre-filtered the dataset to include only ANA positive or negative samples for their corresponding comparisons. In order to improve the fit under a Poisson distribution, CDR3 nucleotide length was transformed to account for the minimum value and step size inherent in the data. An offset variable was created by subtracting 9 (the minimum CDR3 length) from the observed CDR3 nucleotide length and dividing by 3. In order to establish statistical inference, we used the ANOVA function from the car package [44] to obtain an ANOVA-like table, assessing the overall significance of the B cell population. Post-hoc pairwise comparisons between the three levels of Population were performed using the emmeans package. Tukey’s method was applied to adjust for multiple comparisons.

SHM distribution analyses. We performed a Generalized Linear Mixed Models (GLMMs) to analyze the relationship between the percentage of mutated nucleotides and PGA or ANA status of a cell subset. We model the interaction between the factor of interest and B cell populations and added as a random factor the subject of origin to account for clustering caused by the origin of the sample. To ensure a better fit under beta regression SHM values were transformed using the following formula: (% of mutated clones / 100 * (n - 1) + 0.5) / n; where “n” is the total number of observations. To test for statistical significance, we conducted post-hoc pairwise comparisons.

Estimated marginal means were computed, and contrasts were performed to compare levels within each factor. To compare differences for AID targeting to hotspots between ANA+ and ANA-ABCs and memory B cells we performed a pairwise comparison of the average ratio for each gene using Wilcoxon signed ranked test including genes that were present in ANA+ and ANA-subsets. The association between the total number of mutations per cell and the AID targeting ratio was investigated using linear regression. A scatter plot visualizing this relationship, along with fitted linear regression lines for each cell subset, was generated using the ggplot2 package in R. Linear regression with 95% confidence intervals was performed using the lm function, assuming an ordinary least squares model.

Statistical analyses were performed using R version 4.1.1 and the glmmTMB package version 1.1.10 and in graphPad Prism version 10.3.1

## RESULTS

### Clinical data

We included 13 patients, 10 (77%) of whom were female, with a mean ± SD age of 38.3 ± 11.5 years. According to the PGA score, 5 patients were inactive (PGA <0.5), 6 had mild activity (PGA ≥0.5 to 1), and 2 had moderate activity (PGA >1 and ≤2). Patients with mild and moderate activity according to PGA score are considered active. Five patients had a clinical SLEDAI (cSLEDAI) of 0, reflecting inactive disease. Most patients were receiving hydroxychloroquine and different doses of steroids or another immunosuppressive agent. Clinical and laboratory data are provided in Table 1 and in Supplemental Tables 1, 2 and 3.

**Table 1.** Demographic data.

| ID | Age | Sex | Hispanic<br>Ethnicity | Race | Years<br>since<br>diagnosis | SLEDAI-2k | cSLEDAI-2k | PGA | Damage<br>score | Steroid<br>dose* | HCLQ<br>(mg) | Other<br>immunosuppressants | History of<br>Renal<br>Disease |
| --- | --- | --- | --- | --- | --- | --- | --- | --- | --- | --- | --- | --- | --- |
| F292 | 43 | F |  | Black | 18 | 0 | 0 | 0 | 0 | 2 | 400 | - | YES |
| B021 | 42 | F |  | Black | 19 | 10 | 6 | 0.4 | 6 | 3 | 400 | - | YES |
| F049 | 49 | M |  | Black | 13 | 6 | 2 | 2 | 3 | 0 | 0 | Mycophenolate | NO |
| F499 | 22 | M |  | Not identified | 7 | 5 | 0 | 1 | 3 | 30 | 400 | Mycophenolate | YES |
| F135 | 39 | F | Yes | White | 11 | 8 | 4 | 1 | 0 | 0 | 400 | - | NO |
| F487 | 47 | F | Yes | White | 6 | 4 | 2 | 0.2 | 0 | 5 | 400 | Mycophenolate/<br>Belimumab | NO |
| F653 | 23 | F |  | Black | 7 | 0 | 0 | 0.5 | 0 | 40 | 200 | - | NO |
| F335 | 29 | M | Yes | White | 11 | 8 | 4 | 1.5 | 0 | 0 | 400 | - | NO |
| F139 | 59 | F |  | Black | 23 | 2 | 2 | 0.8 | 0 | 0 | 400 | - | NO |
| F155 | 36 | F |  | Black | 11 | 2 | 0 | 0.4 | 2 | 0 | 400 | Azathioprine | YES |
| F326 | 25 | F |  | Black | 8 | 2 | 0 | 0.3 | 0 | 2 | 400 | - | NO |
| F104 | 34 | F | Yes | Not identified | 21 | 6 | 4 | 1 | 0 | 3 | 300 | - | YES |
| F124 | 50 | F |  | Not identified | 26 | 7 | 2 | 1 | 0 | 2 | 400 | Mycophenolate | YES |

\*(equivalent mg of prednisone per day)
F: Female. M: Male. SLEDAI: Systemic lupus erythematosus disease activity index. cSLEDAI: Clinical systemic lupus erythematosus disease activity index. PGA: Physician Global Assessment. HCLQ: hydroxychloroquine.

### B cell immunophenotyping

There was considerable variability in the percent of each subset among circulating B cells across patients (Figure 1C and 1D). As expected, naïve B cells were the most abundantly represented subset with a median of 44% of circulating B cells classified as naïve. The median percentage of ABCs within total B cells was 6%. In all but 2 patients, the percentage of circulating plasmablasts was <5%; in the two patients with higher plasmablast fractions, this subset was considerably expanded representing 11% and 53% of the B cells (Figure 1C). A weak correlation between frequency of ABCs and plasmablasts in patients with SLE was previously reported [30]; however, we did not confirm this association (supplemental Figure 1A). We also investigated the association between the percent of plasmablasts and other B cell subsets and found a significant negative correlation between the percent of plasmablasts and naïve cells (r=-0.632, R^2^=0.400, p=0.02) (Supplemental Figure 1B). There was correlation between the percent of switched and unswitched memory B cells (r=0.58, R2=0.31, p=0.049) (Supplemental Figure 1C). Consistent with previous reports, most of the ABCs were class switched (IgD-) (Supplemental Figure 1D). Next, we analyzed the subset composition of ANA+ B cells. A median of 8.3% (IQR 6-9.8%) of naïve B cells were ANA+ in this cohort (Figure 1E). These results are in line with previous studies which found that a relatively high percentage of naïve B cells are autoreactive in both healthy individuals and individuals with SLE [9,5,4]. An immune checkpoint that reduces the percentage of autoreactive naïve B cells transitioning to the memory B cell subset has been reported previously [45–47,4,48]. The percent of ANA+ B cells was similar among naïve B cells and ABCs (median 8.3%, IQR 4.8-11.9%) [14,30]. ANA positivity was significantly higher in both naïve B cells and ABCs than in memory B cells (median 4.1%, IQR 3.3-6.3; p<0.05 both comparisons) (Figure 1E).

### B cell receptor repertoire sequencing

To gain insights into the repertoire features of the ANA+ vs. ANA-B cells, antibody heavy chain gene rearrangements were amplified from bulk sorted populations (see methods). Total clone counts per donor for ANA+ and ANA-ranged from 97,860 in F292 to 5460 in F499 (Figure 1F). In nearly all cases, we observed more B cell clones in ANA-compared to the ANA+ B cells, consistent with the greater presence of ANA-B cells from each subset during the sorting procedure (Figure 1G). Furthermore, in most individuals, the percentage of clones from naïve cells was higher than memory cells or ABCs (Figure 1H), consistent with the flow cytometry data and general findings about diversity in the naïve B cell pool [49]. The percentage of sequences from each individual obtained from plasmablasts was highly variable (Supplemental Table 4). There was no association between the number of clones and disease activity (Supplemental Figure 1E).

To determine if ANA+ BCRs had different repertoire features from ANA-BCRs, we analyzed clone size distributions, VH gene usage, CDR3 length distributions, and SHM frequencies and distributions. To determine if some of the samples were dominated by very large clones, we computed the D20 index, which represents the sum of the copy numbers for the top 20 ranked clones in each sample divided by the total copies in the sequencing library (Supplemental Figure 1F and 1G). This analysis revealed considerable heterogeneity. We observed an apparent bias towards larger clone sizes in ANA+ vs. ANA-cells, with more ANA+ clones belonging to the top 20 clones (Supplemental Figure 1F), however, D20 was highly correlated with differences in the total clone count per sorted sample (Supplemental Figure 1G) and may not necessarily reflect the underlying clonal architecture.

### VH and JH usage and CDR3 length in B cell subsets

We analyzed VH and JH gene usage, aggregating the data for each subset per individual donor (Figure 2A and 2B). Although there is intra-donor variability and not all IGHV genes are equally represented across donors, six IGHV genes were most frequently used by the patients in the study: IGHV1-18, IGHV3-21, IGHV3-23|3-23D, IGHV4-34, IGHV4-39 and IGHV4-59 (Figure 2A). We observed that some genes were preferentially used by ANA+ or ANA-B cells (Figure 2A). We compared VH gene usage between ANA+ and ANA-B cells across different subsets for the top 6 expressed VH genes (Figure 2C), IGHV3-21 and IGHV4-39 were preferentially used by ANA+cells in naïve and ABC subsets. IGHV1-18 and IGHV4-59 were preferentially used by ANA+ naïve B cells.

**FIGURE 2.**
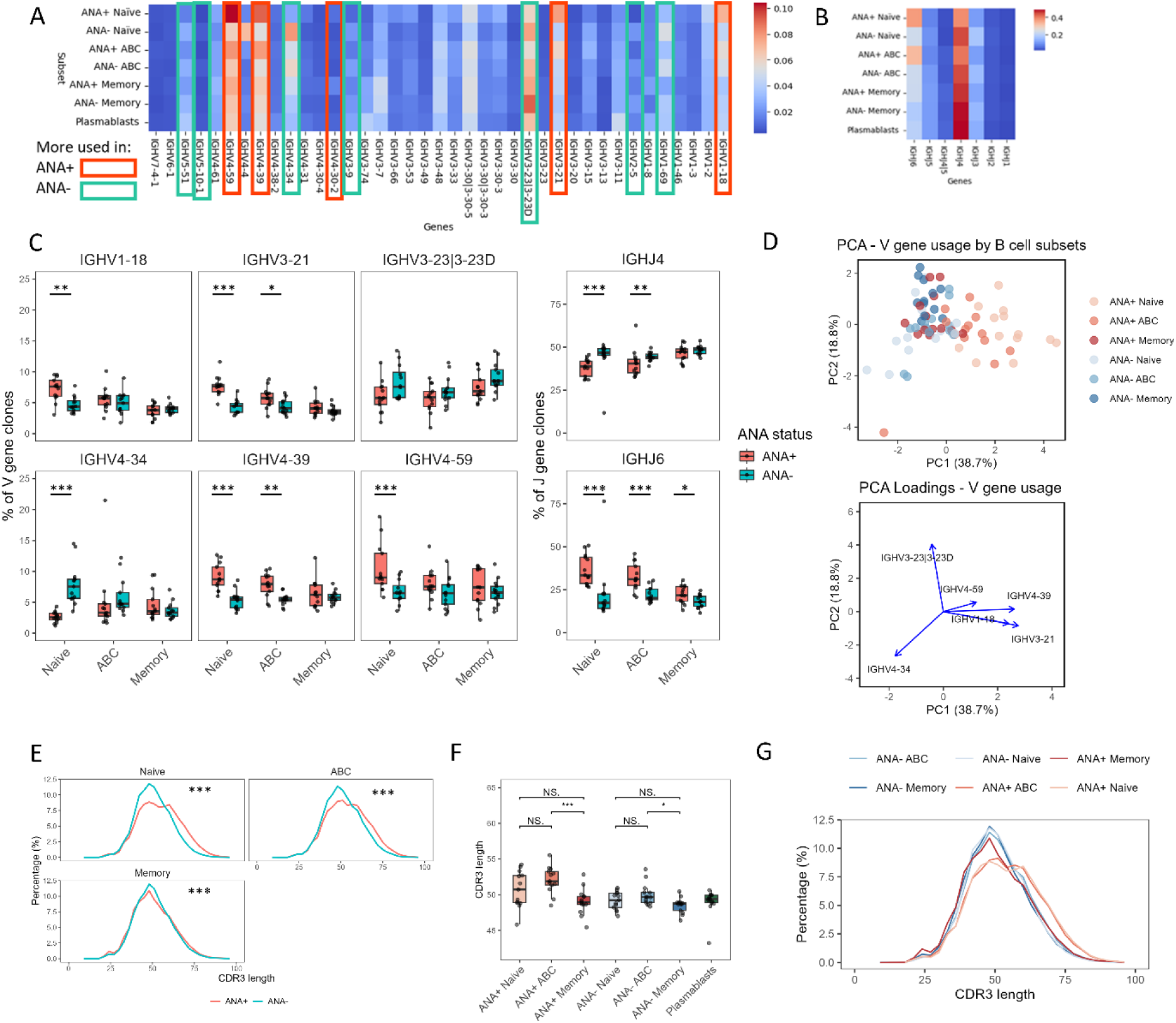
IGHV/J usage and CDR3 length analysis in B cell subsets. A and B) Heatmap of relative IGHV/J gene usage by subset. Data are based on clone counts (each clone is counted once). Results are normalized by row. Genes present in <0.5% of clones were excluded. The color scale indicates normalized clone count fraction. In A) red rectangles highlight gene segments that are more frequently used by ANA+ B cells and green those more frequent on ANA-B cells. Comparisons were performed using Wilcoxon rank sum test with FDR correction; rectangles indicate significant differences. C) Boxplots represent the proportion of ANA+ and ANA-clones utilizing the six most prevalent VH and two most prevalent JH genes, divided by B cell subsets. Each dot represents an individual patient. Comparisons were performed between each ANA+ and ANA-pair using Wilcoxon signed rank test with FDR correction. D) Principal component analysis of the six most prevalent VH genes in each B cell subset, by donor. Data were scaled and centered before PCA. E) Histogram of the direct comparison of CDR3 length distribution for ANA+ vs ANA-subsets. Comparisons were performed using a GLMM as described in methods. F) Boxplots show the mean CDR3 length for each subset; each dot represents an individual patient. Pairwise comparisons between ANA+ subsets and ANA-subsets were performed with Wilcoxon signed rank test with FDR correction. G) Histogram showing the distribution of CDR3 lengths for each subset. Graphs show comparisons that were significant after FDR correction. Adjusted p value: *p < 0.05, **p < 0.01, ***p < 0.001. ANA: Antinuclear antigens. FDR: False discovery rate. GLMM: Generalized linear mixed model.

To investigate similar gene usage by different ANA+ and ANA-B cell subsets, we performed a principal component analysis including the 6 more prevalent genes in ANA+ and ANA-negative subsets. We observed that ANA+ ABC and ANA+ naïve B cells cluster together, separated from ANA-subsets and ANA+ memory B cells (Figure 2D). This discrimination was mainly driven by IGHV3-21, IGHV4-39 and IGHV1-18 in PC1. This suggests that there is selection against genes that are frequently used in ANA+ naïve B cells in memory B cells, and that ANA+ naïve B cells can differentiate into ANA+ ABCs without being subject to this tolerance checkpoint.

The gene IGHV4-34 is of special interest. This gene is overrepresented in the repertoire of patients with SLE [50] and it has been associated with clinical manifestations of SLE such as nephritis and neuropsychiatric involvement [51]. One specificity of antibodies produced by cloned sequences that use this gene is the I/i antigen of red blood cells [50]. Antibodies encoded by IGHV 4-34 also can bind single-stranded (ss)DNA, lipid A, cardiolipin, rheumatoid factor and CD45, a cell surface protein [51–53]. A subset of the antibodies produced by IGHV4-34 with the 9G4 idiotype, characterized by conserved germline AVY and QW amino acid motifs in framework-1 region, are associated with SLE severity [21]. We found that the IGHV4-34 gene was more frequently used in ANA-naïve B cells (Figure 2B).

We also explored IGHJ gene usage by different B cell subsets (Figure 2B). IGHJ6 was more frequently used by all ANA+ subsets and IGHJ4 by ANA-naïve cells and ABCs (Figure 2C). IGHJ4 was also more prevalent in plasmablasts, suggesting preferential differentiation of ANA-cells to the plasmablast pool. We did not observe differences in VH or JH gene usage between patients with active or inactive disease (Supplemental Figures 2A-C).

CDR3 length distribution differences are a signal of repertoire variations among individuals and over time [54]. Long CDR3 sequences have been associated with autoreactivity in early B cells and selection against antibodies with long CDR3 regions has been described [55]. We observed longer mean CDR3s in ANA+ naïve B cells and ABCs when compared with their ANA-counterparts (Figure 2F). In memory B cells ANA+ cells have a slightly shorter CDR3 compared to ANA-memory B cells (Figure 2E). Mean CDR3 length from ANA+ and ANA-ABCs was longer than memory B cells (Figure 2F). Overall, ANA+ naïve and ANA+ ABCs have longer CDR3 sequences (Figure 2G). There was no difference in CDR3 length according to disease activity status (Supplemental Figure 2D). Thus, these data also suggest that ANA+ naïve B cells with long CDR3 sequences are negatively selected as they transition to become memory cells and that this selection does not affect the transition to the ANA+ ABC compartment.

### Somatic hypermutation in B cell subsets

As expected, there was minimal SHM in naïve B cells (Figure 3A). ABCs exhibited SHM supporting antigen experience (Figure 3A). We observed a higher frequency of SHM in both ANA+ and ANA-memory B cells compared to ANA+ and ANA-ABCs (Figure 3B). The frequency of SHM in memory B cells, especially in ANA-memory B cells, was more similar to what we observed in plasmablasts (Figure 3B and Supplemental Figure 3A). Thus, the frequency of SHM is highest in plasmablasts and memory B cells, with a lower frequency in ABCs and almost no SHM in naive B cells. In both ABCs and memory B cells, ANA+ cells exhibited less SHM compared to ANA-cells (Figure 3C); the difference was more pronounced in ABCs. There was a small difference between ANA+ vs ANA-naïve B cells that was statistically significant, but probably not biologically relevant (Figure 3C). The difference observed in SHM between ANA+ and ANA-B cells when all genes combined were analyzed, was also observed in specific genes of interest, such as IGHV4-34 and IGHV4-59 (Supplemental Figure 3B). Patients with active disease had lower levels of SHM in ABCs, both ANA+ and ANA-, and in ANA+ memory B cells (Figure 3D).

**FIGURE 3.**
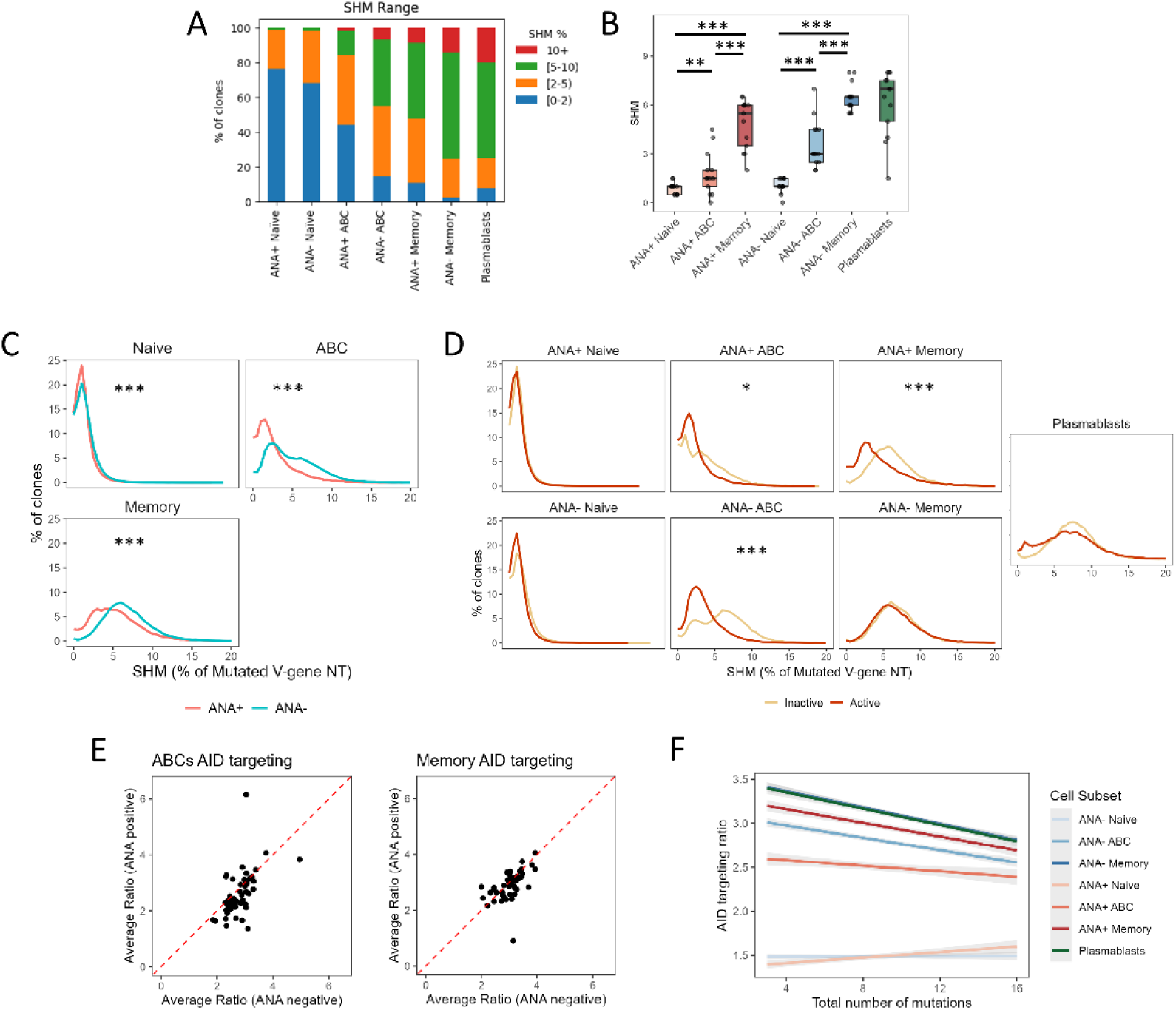
Frequency of SHM in B cell subsets. A) Frequency of clones with different ranges of SHM for each subset. SHM was calculated from pooled clones across all donors for each subset. B) Median value per individual patient of the percent of mutated V-gene nucleotides for each subset. Pairwise comparisons between subsets within ANA+ ANA-subsets were performed using Wilcoxon signed rank test with FDR correction. C and D) Histograms of the direct comparison of clones with mutated V-gene nucleotides between: ANA+ and ANA-subsets (C) or between active and inactive patients according to PGA (D). Comparisons were performed using a GLMM as described in methods. Graphs show comparisons that were significant after FDR correction. Adjusted p value: *p < 0.05, **p < 0.01, ***p < 0.001. E) Scatterplot of the average ratio (observed/expected) of AID targeting of ANA+ vs ANA-ABCs and memory B cells. Each doth represents the average of an individual IGHV gene. Wilcoxon signed-rank test, p-value for ABCs <0.001, for memory B cells p=0.03. F) Scatter plot of the linear association between total number of mutations and AID targeting ratio for each cell subset using linear regression. The shaded area around each regression line represents 95% confidence interval. SHM: Somatic hypermutation. ANA: Antinuclear antigens. FDR: False discovery rate. PGA: Physician Global Assessment. GLMM: Generalized linear mixed model.

### AID targeting to hotspots

To evaluate AID targeting to hotspots, this calculation was performed dividing the observed mutation frequency at WRC sites (number of mutations at WRC sites / total number of mutations) by the expected frequency of WRC sites (number of WRC sites / sequence length). This enrichment ratio was then compared between ANA+ and ANA-ABCs and memory B cells. We did not include naïve B cells because of the low mutation frequency in this subset. We observed a higher AID targeting ratio in genes derived from ANA-ABCs and, to a lesser extent, memory B cells (Figure 3E). This suggests increased AID activity and/or selection for mutations at WRC hotspots in ANA-subsets. Given the higher overall mutation frequency observed in ANA-cells, we investigated whether the differences in AID targeting ratios could be attributed simply to differences in mutation ratio. To address this, we analyzed the relationship between the total number of mutations and the AID targeting ratio for each B cell subset (Figure 3F). We observed a gradual decrease in the AID targeting ratio as the total number of mutations increased in ABCs, memory B cells, and plasmablasts. The regression lines for ANA-memory B cells and plasmablasts largely overlapped. Importantly, the differences between ANA+ and ANA-ABCs and memory B cells in AID targeting ratios, and their 95% confidence intervals, remained distinct across the range of mutation frequencies analyzed. This supports the conclusion that the observed differences in AID targeting are not only a consequence of the different mutation frequencies between these subsets.

### Clonal association with plasmablasts

We explored clonal relatedness by quantifying clones with members in multiple subsets. We observed that overall, most sequences were unique and not present in other subsets (Figure 4A). We observed that memory B cells share more sequences with plasmablasts, followed by ABCs. Naïve B cells shared less sequences, likely consistent with higher diversity and smaller clone size in the naïve B cell pool (Figure 4). More ANA-ABCs and ANA-memory B cells shared sequences with plasmablasts than ANA+ B cells (0.75% vs 0.3%, p<0.001) or ANA+ memory B cells (2.85% vs 1.23%, p<0.001). The opposite was observed in naïve B cells in which more ANA+ B cells shared sequences with plasmablasts (0.12% vs 0.07%, p=0.02), perhaps suggesting that these cells differentiated through an EF route.

**FIGURE 4.**
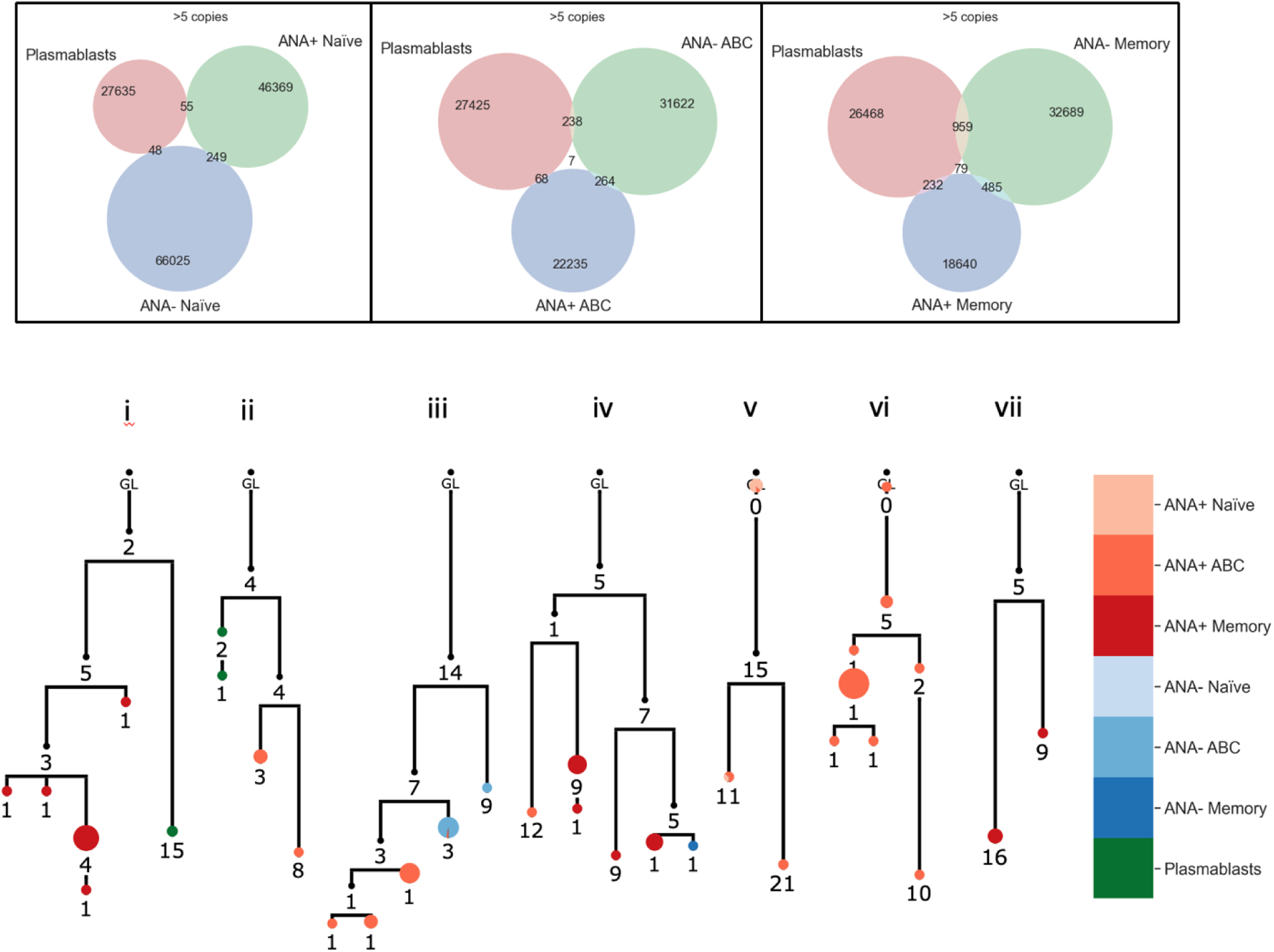
Clonal relatedness of plasmablasts sequences. A) Venn diagram shows the number of individual clones that were identified for each subset and the number of clones shared between subsets. Only clones with more than 5 copies were included. B) ANA+ Clonal lineages exhibit different subset distribution patterns. i, ii) separate branching in different subsets; iii, iv) ANA+ and ANA- in the same lineage; v) shared unique sequences; vi, vii) predominantly single subset. Nodes are colored by subset and proportional to copy number. Nodes with wedges contain more than one subset. Black nodes are inferred. Numbers refer to mutations in VH compared to preceding vertical node. Clone metadata-donor, clone ID, VH gene and CDR3 amino acid sequence: i) F049_478961_IGHV3-30-3_CARGNYYYGSGSYSLDYW; ii) F104_713843_IGHV3-21_CARESERYYYDTSGYSPMDYW iii) F049 411445 IGHV1-8 CARRFYYDNGGYSGTYDYW; iv) F135_1194397_IGHV3-64_CARGGYDSSGTVGLKNWYFDLW; v) F139_1463574_IGHV3-7_CARKGYYYDSSGYYNTFDYW; vi) F124_845512_IGHV1-8_CARRRTWGITPYYFDSW; vii) F049_510114_IGHV3-30|3-30-5_CAKDPQYYDSWSGYSGEVYYYYYNMDVW

The association between SHM and autoreactive BCRs in antigen experienced cells at the single clone level is complex. There are data that support that autoreactivity in memory B cells can be generated de novo by SHM of ANA-B cells; however, SHM can also modify the specificity of autoreactive B cells, generating non-autoreactive clones [45,56–59]. Lineage analysis revealed clones with all members ANA+, ANA- or both ANA+ and ANA-members of the same clone (Figure 4B). The presence of clones in which individual cells have a different ANA status highlights the fact that SHM can change antigenic specificity. It could also represent pairing with different light chains, which we did not interrogate.

## DISCUSSION

To get some insight into the derivation of autoreactive B cells in patients with SLE, we performed BCR heavy chain sequencing of different B cell subsets: naïve, ABCs, and memory B cells, subsetted based on their ANA reactivity; and plasmablasts.

We observed enrichment of some IGHV genes in different B cell subsets. The enrichment of IGHV3-21and IGHV4-39 in ANA+ naïve, ANA+ ABCs, but not in memory subsets could suggest that this gene is prone to autoreactivity and extrafollicular differentiation. Over-utilization of IGHV3-21 has been reported in mantle cell lymphoma, where clones often exhibit SHM and lambda light chain restriction, which could indicate antigen selection [60]. IGHV4 family members, including IGHV4-39 and IGHV4-59, have been found at increased frequencies in cerebrospinal fluid and in central nervous system from patients with multiple sclerosis [61–63], but their ties to autoreactivity are unclear. We also observed that IGHV1-18 contributed to the clustering of naïve ANA+ and ABC ANA+ compartments separate from other B cell subsets, but it was only more frequent in ANA+ compared to ANA-naïve B cells. In patients with acute dengue infection positive selection of IGHV1-18 expressing B cells has been reported and it is speculated that these cells differentiate through an EF pathway [64].

Finally, we observed that IGHV4-34 is enriched in the ANA-naïve B cell subset of patients with SLE. It is not surprising that VH4-34 is frequently encountered in naïve B cells, including naïve B cells from SLE patients [22]. Considering that this gene is associated with autoreactivity, it seems counterintuitive that many of the naïve VH4-34 B cells are ANA-; however, many of the antibodies encoded by this gene can react with RBC and B cells [50], and not necessarily with nuclear antigens [65]. IGHV4-34 was frequently used by ABCs, and this is consistent with previous reports of an expansion of 9G4+ B cells within ABC subsets: activated naïve (IgM+) [22] and DN2 (class switched) B cells [14]. It was also previously reported that IGHV4-34 is negatively selected in the germinal center and the memory B cell compartment [66]. We also found that IGHV4-34 has a low prevalence in the memory compartment.

With respect to overall IGHV usage, ANA+ ABCs, are more similar to ANA+ naïve B cells than to memory B cells according to principal component analysis. This suggests a direct differentiation process, which is not subject to the same immune checkpoint that precludes the differentiation of ANA+ naïve B cells into memory B cells.

We did not include healthy controls to compare the relative frequency of clones using different IGHV genes; however, there are previous studies that analyzed BCR sequences in patients with SLE compared to healthy individuals. A preferential usage of IGHV4 genes and IGHV1-18 in patients with SLE was previously reported by Bashford-Rogers [67]. In a case series of patients with anti-N-methyl-D-aspartate receptor (NMDAR) encephalitis (an autoantibody-mediated disorder) IGHV1-18 was also associated with disease [68]. Ota et al. analyzed BCR gene expression in B cell subsets and compared this expression with healthy donors [20]. They found increased usage of gene IGHV4-59 across multiple subsets in patients with SLE; they also found increased expression of IGHV1-18 and IGHV3-21 in unswitched memory B cells from patients with SLE [69]. The increased frequency of these IGHV genes in SLE patients suggests their potential contribution to pathogenesis, a finding further supported by our observation that these genes are more frequent in ANA+ subsets.

We also observed a preferential use of two IGHJ segments, IGHJ4 and IGHJ6, similar to findings by others [70]. IGHJ4 was preferentially used by all subsets, but more frequent in ANA-subsets, while IGHJ6 was more frequent in ANA+ subsets, suggesting a role in generation of autoreactivity. This is consistent with previous reports showing a progressively increased use of IGHJ4 and diminished use of IGHJ6 as B cells differentiate from transitional to naïve and memory compartments [71]. IGHJ6 encodes a long stretch of tyrosine residues that contribute to increased CDR3 length [72]. Our data suggest that the progressive reduction in IGHJ6 usage correlates with loss of longer CDR3 sequences [48] and selection of non-autoreactive clones.

Longer CDR3s were previously reported in naïve B cells from healthy individuals when compared with plasmablasts, suggesting a preferential selection of short CDR3 segments into the plasmablast compartment [20]. However, those analyses did not identify autoreactive cells.

We observed that plasmablasts have a CDR3 length similar to ANA-naïve B cells and shorter CDR3 sequences than ANA+ naïve B cells, supporting a preferential differentiation into plasmablasts of B cells with a shorter CDR3 and the presence of an immune checkpoint that reduces the differentiation of ANA+ naïve B cells with long autoreactive CDR3 segments in the plasmablast compartment.

We also observed that ANA+ memory B cells have CDR3 sequences that are more similar to ANA-naïve B cells, and shorter than ANA+ naïve B cells, suggesting that many ANA+ memory cells may have originated from ANA-precursors and acquired autoreactivity by SHM. This is consistent with previous studies that showed that autoreactive sequences obtained from memory IgG+ cells could be created de novo by SHM [45]. In contrast to what we observed in memory B cells, ANA+ ABCs have longer CDR3s, similar to ANA+ naïve cells, and longer than the CDR3 of ANA-ABCs, which might suggest that ANA+ ABCs arise through an EF pathway from ANA+ naïve B cells and undergo a different selection process than conventional memory B cells. SHM has been traditionally associated with GCs. Although it has been reported that SHM can occur outside of the GC [73], it is expected that cells generated through EF reactions will have a lower frequency of mutations [14,74]. We observed that ABCs are somatically hypermutated, which is in line with reports that suggest that ABCs represent atypical memory B cells [75] and consistent with previous reports that specifically showed SHM in this subset [76]. The fact that memory B cells have a higher frequency of SHM compared to ABCs is consistent with an EF origin of ABCs.

We observed less SHM in ANA+ ABCs and memory B cells compared to ANA-subsets. Considering that SHM is associated with selection and affinity maturation [55], our results suggest that antigen experienced ANA+ B cells, and especially ANA+ ABCs are subject to less stringent selection. This may reflect either an early exit from GCs or EF differentiation. ANA+ and ANA-ABCs and ANA+ memory B cells showed lower frequencies of SHM in patients with active disease, suggesting that the regulation of SHM is a dynamic process and potentially more cells are being generated through EF pathways in patients with active disease. We also observed a relatively high ratio of SHM in plasmablasts, consistent with previous studies of plasma cells obtained from the bone marrow in healthy individuals which showed a significant increase in the frequency of SHM compared with circulating memory B cells [77]. Considering the dispersion of the data, it is possible that plasmablasts in SLE have a mixed origin, some of them coming from EF and others from GC responses [14], with EF generated plasmablasts showing less SHM than GC generated plasmablasts [74].

There is evidence of AID expression [78] and class switch recombination, which is also dependent on AID, outside of the GC [79]. Additionally, SHM has been described out of the GC [73]. However, it is generally accepted that most of the SHM associated with affinity maturation occurs in the GC. Thus, it is possible that increased AID targeting reflects a preferential GC origin of some subsets. To gain more insight into the selection process of ANA+ and ANA-subsets, we analyzed mutation targeting to hotspots. We observed that AID appears to be mutating specific hotspot regions within antibody genes (WRC hotspots) in ANA-subsets in ABCs and memory B cells. This could mean that AID is either more active in these cells or that there is stronger selection pressure favoring mutations at these specific hotspots. The fact that among the antigen experienced cells, ANA+ ABCs showed the lowest AID targeting ratio, might support an EF origin of these cells.

It has been proposed that ABCs are relevant to SLE pathogenesis. There is a lack of a clear definition in the literature about the phenotype or subsets of ABCs. Our definition includes activated naïve [22], DN2 [14] and CD11c+ [30] B cells. Although their function has not been clearly established, it has been proposed that ABCs can function as antigen presenting cells, or they can differentiate into antibody secreting cells. We found a relatively low number of sequences shared between memory B cells and plasmablasts, and even fewer between ABCs and plasmablasts. In both cases, there were more shared clones in ANA-than in ANA+ subsets. This supports the presence of an immune checkpoint that prevents the differentiation of ANA+ ABCs into plasmablasts. We found evidence of SHM in ABCs that supports that they are antigen experienced. The frequency of autoreactivity in ABCs is more than in memory B cells and similar to naïve B cells, suggesting that ABCs and memory B cells are not subject to the same immune checkpoint, and highlighting the potential relevance of ABCs for the generation of auto-antibodies. Furthermore, differences in gene segment usage, CDR3 length, and SHM rate support a close relationship between ANA+ ABCs and ANA+ naïve B cells, and that the evolution of autoreactive ABCs is different from that of memory B cells. Overall, the results of our analyses of ANA+ B cells suggest that ANA+ ABCs are more similar to ANA+ naïve, and distinct from memory B cells.

Our study has some limitations: We analyzed mixed memory B cell subsets (IgM and IgG for memory and ABCs), and limited our analysis to heavy chains without analyzing light chains. We studied a limited number of patients and we only looked at a single time point per patient, which prevented us from evaluating changes associated with disease flare or quiescence. Finally, we were not able to explore the ANA reactivity of plasmablasts because of technical limitations, but we were able to evaluate clonal overlap between plasmablasts and other B cell subsets.

## CONCLUSIONS

Based on BCR sequences, ANA+ ABCs are similar to ANA+ naïve B cells, and exhibit distinct features compared to memory B cells, including longer CDR3 and differential IGHV gene usage. This suggests a different selection process for autoreactive ABCs, bypassing an immune checkpoint that prevents differentiation of ANA+ naïve B cells into memory cells.

ANA+ ABCs and ANA+ memory B cells have relatively lower rates of somatic hypermutation than ANA-cells, and different AID targeting to mutational hotspots, suggesting an EF origin and supporting that the EF pathway is important for generation of autoreactivity. Patients with active SLE have evidence of increased EF differentiation of B cells.

