## Supplemental figures 1 - 3 for "SELECTION OF ANTI-NUCLEAR ANTIGEN (ANA) REACTIVE B CELLS IN SYSTEMIC LUPUS ERYTHEMATOSUS"

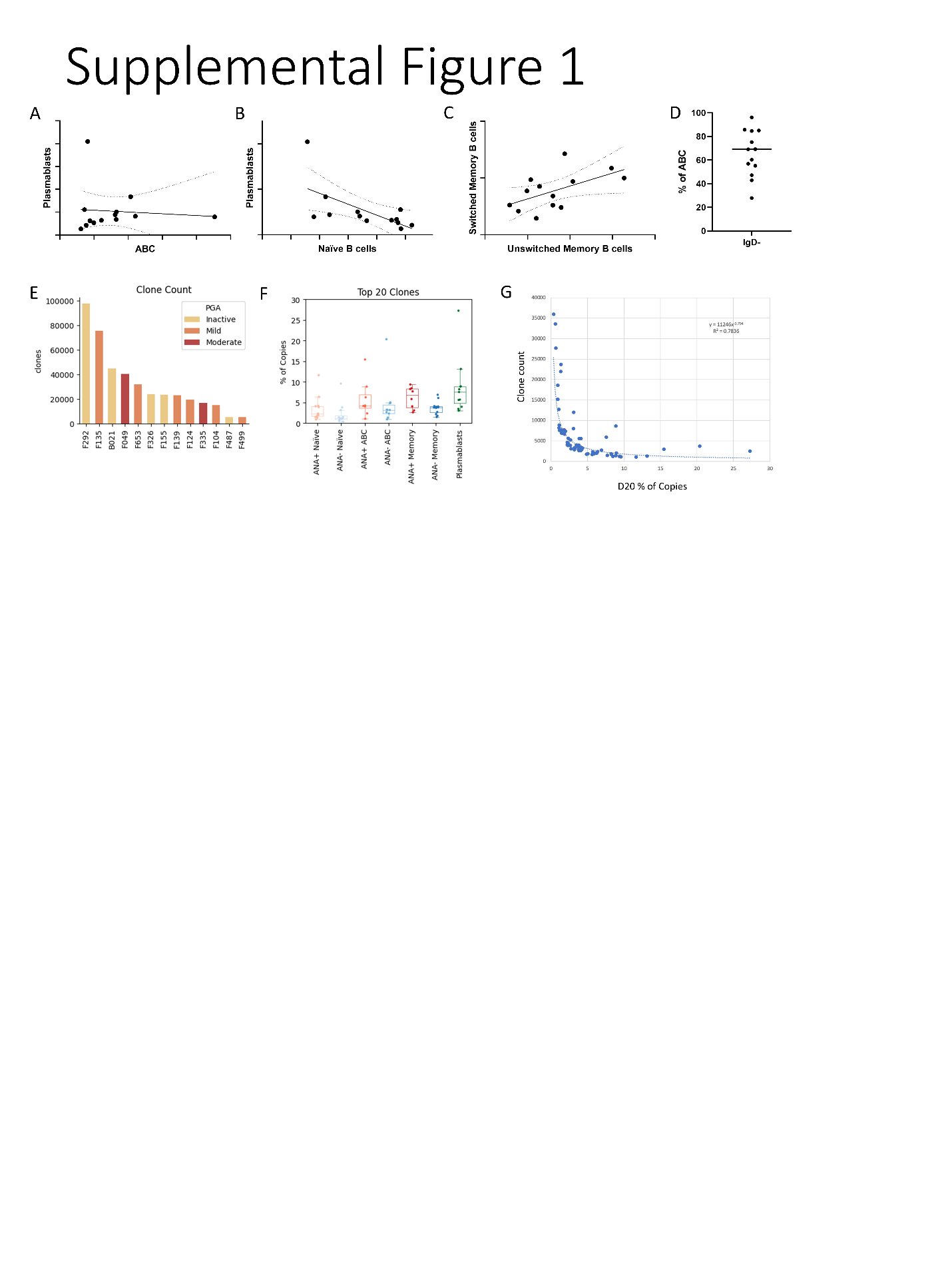

SUPPLEMENTAL FIGURE 1. A, B and C) Correlation of the percentage within B cells of different B cell subsets. To evaluate linear associations, we performed arcsine transformation of the raw percentages. The graphs show a regression line with 95% confidence intervals. *p<0.05. D) Percentage of IgD- cells (class switched) within the ABC subset. Each dot represents an individual patient. E) Histogram of number of clones identified in each patient. Columns are colored according to disease activity status (inactive = PGA <0.5; mild activity = PGA ≥0.5 to 1; moderate activity = PGA >1 and ≤2). F) Boxplots of the percentage of copies represented by the top 20 most abundant clones by subset. Each dot represents an individual patient. G) Scatterplot representing the association of the percentage of copies represented by the top 20 most abundant clones in each subset for all donors with the total number of clones in each subset. PGA: Physician Global Assessment

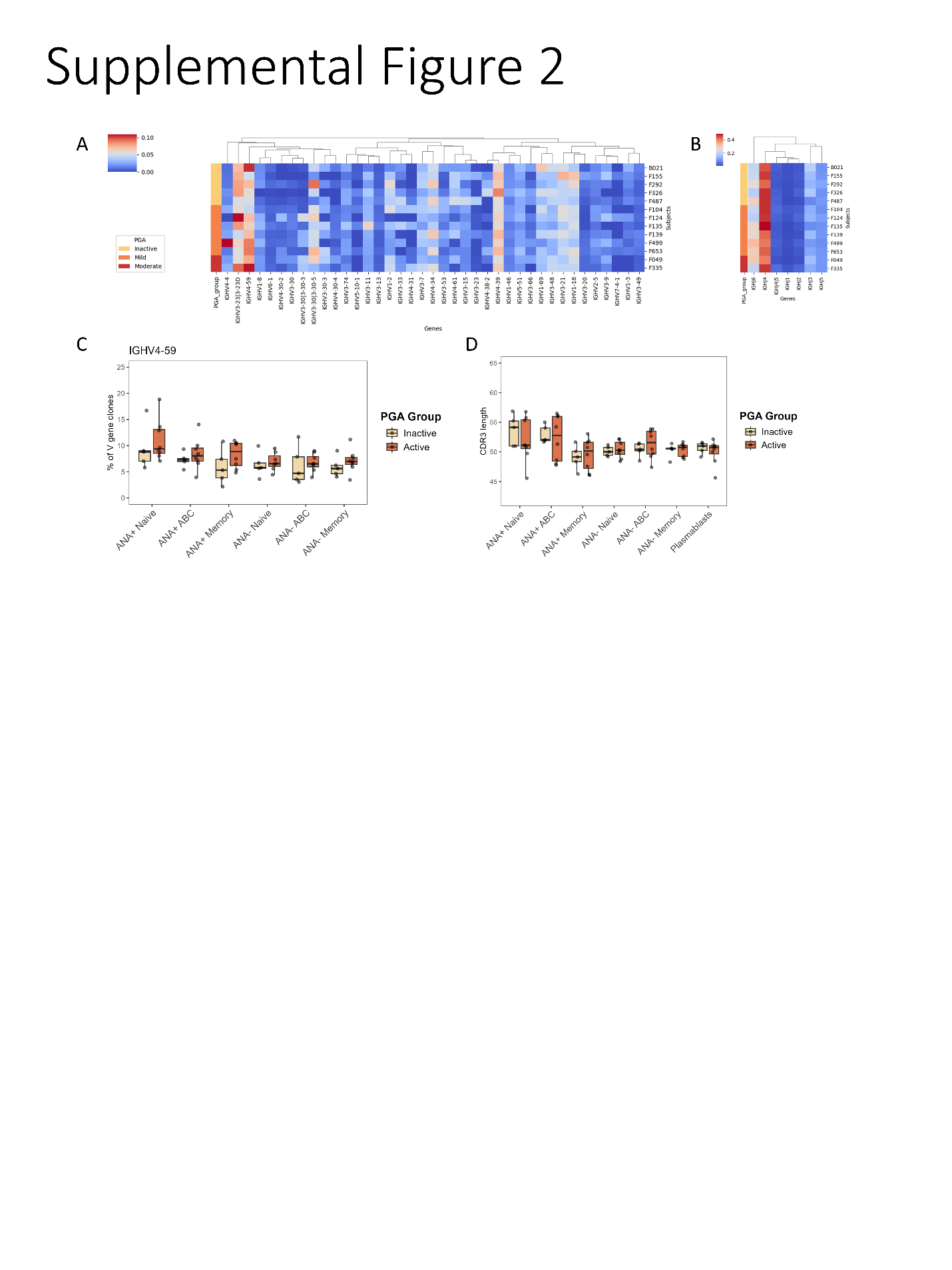

SUPPLEMENTAL FIGURE 2. A and B) Heatmap of relative IGHV/J gene usage by subset. Data are based on clone counts (each clone is counted once). Results are normalized by row. Genes present in <0.5% of clones were excluded. The color scale indicates normalized clone count fraction. Left color scale represents disease status group according to PGA (inactive = PGA <0.5; mild activity = PGA ≥0.5 to 1; moderate activity = PGA >1 and ≤2). C) Boxplots of the proportion of IGHV clones utilizing IGHV4-59 divided by B cell subsets and stratified by disease activity status according to PGA (inactive = PGA <0.5; active = PGA ≥0.5). Each dot represents an individual patient. D) Boxplots show the mean CDR3 length for each subset divided in active e inactive patients as described in C. Each dot represents an individual patient. In panels C and D comparisons were performed using Wilcoxon rank sum test (non-paired); no significant differences were observed.

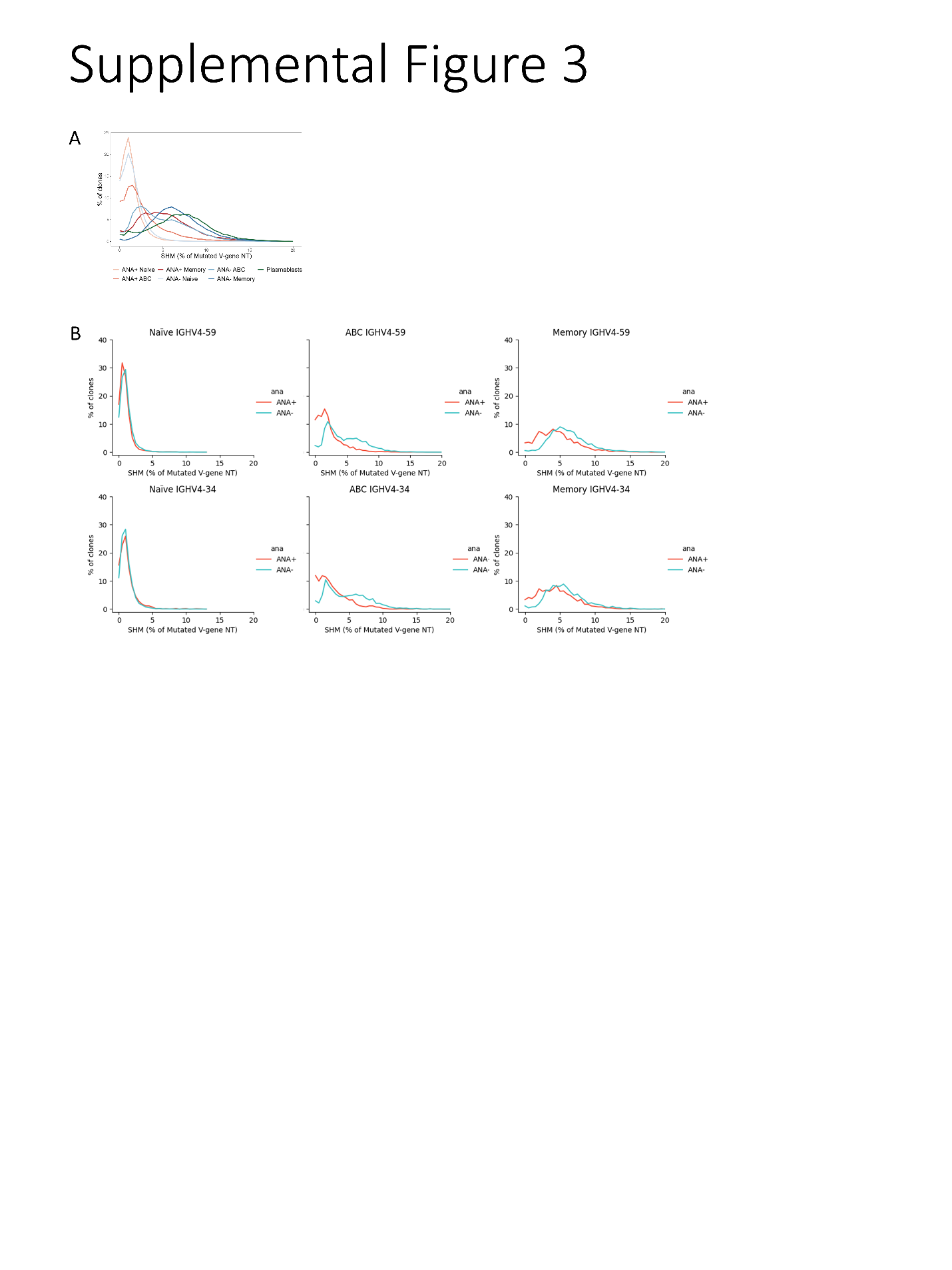

SUPPLEMENTAL FIGURE 3. A) Histogram showing the percentage of clones according to their frequency of mutated V-gene nucleotides for each B cell subset. B) Histograms of the direct comparison of clones with mutated V-gene nucleotides between ANA+ and ANA- subsets for IGHV4-59 and IGHV4-34.
