## Supplemental tables 1 - 3 for "SELECTION OF ANTI-NUCLEAR ANTIGEN (ANA) REACTIVE B CELLS IN SYSTEMIC LUPUS ERYTHEMATOSUS"

**Supplemental table 1. Detailed clinical data**

| ID | Rash | Nasal ulcer | Oral Ulcer | Alopecia | CNS | Serositis | Arthritis | Proteinuria | Pyuria | Hematuria | Urinary Cast | Hematologic  cytopenia |
| --- | --- | --- | --- | --- | --- | --- | --- | --- | --- | --- | --- | --- |
| F292 |  |  |  |  |  |  |  |  |  |  |  |  |
| B021 |  | Yes |  |  |  |  |  | Yes |  |  |  |  |
| F049 | Yes |  |  |  |  |  |  |  |  |  |  |  |
| F499 |  |  |  |  |  |  |  |  |  |  |  |  |
| F135 |  |  | Yes | Yes |  |  |  |  |  |  |  |  |
| F487 |  |  |  | Yes |  |  |  |  |  |  |  |  |
| F653 |  |  |  |  |  |  |  |  |  |  |  |  |
| F335 | Yes |  |  |  |  |  |  |  |  |  |  |  |
| F139 | Yes |  |  |  |  |  |  |  |  |  |  |  |
| F155 |  |  |  |  |  |  |  |  |  |  |  |  |
| F326 |  |  |  |  |  |  |  |  |  |  |  |  |
| F104 | Yes |  | Yes | Yes |  |  |  |  |  |  |  |  |
| F124 | Yes |  |  |  |  |  |  |  |  |  |  |  |

CNS: Central nervous system

**Supplemental Table 2. Detailed laboratory data**

| ID | WBC | Platelets | Lymphocyte count | Lymphocyte % | Neutrophil count | Neutrophil % |
| --- | --- | --- | --- | --- | --- | --- |
| F292 | 5.58 | 243 | 1.81 | 32.4% | 3.18 | 57.0% |
| B021 | 3.8 | 139 | 0.6 | 15.8% | 2.6 | 68.4% |
| F049 | 4.18 | 308 | 0.44 | 10.5% | 3.13 | 74.9% |
| F499 | 9.4 | 86 | 0.57 | 6.1% | 7.92 | 84.3% |
| F135 | 4.6 | 323 | 1.1 | 23.9% | 2.96 | 64.3% |
| F487 | 3.94 | 298 | 0.99 | 25.1% | 2.2 | 55.8% |
| F653 | 7.72 | 206 | 1.52 | 19.7% | 5.76 | 74.6% |
| F335 | 7.55 | 278 | 0.5 | 6.6% | 6.59 | 87.3% |
| F139 | 3.4 | 171 | 1.19 | 35.0% | 1.78 | 52.4% |
| F155 | 4.1 | 272 | 1.21 | 29.5% | 2.65 | 64.6% |
| F326 | 4.11 | 268 | 1.14 | 27.7% | 1.63 | 39.7% |
| F104 | 3.24 | 219 | 0.91 | 28.1% | 1.47 | 45.4% |
| F124 | 2.88 | 216 | 1.09 | 37.8% | 1.28 | 44.4% |
| WBC: White blood cells. | | | | | | |

**Supplemental Table 3. Complement levels and autoantibody profile**

| ID | dsDNA | Low  complement | Sm | RNP | Ro | La | ACL IgG | ACLIgA | ACLIgM | LAC | B2GPI IgG | B2GPI IgA | B2GPI IgM |
| --- | --- | --- | --- | --- | --- | --- | --- | --- | --- | --- | --- | --- | --- |
| B021 | + | + | - | - | - | - | - | - | - | - | NA | NA | NA |
| F049 | + | + | + | + | + | - | - | - | - | - | - | - | - |
| F104 | + | + | + | + | + | + | - | - | - | - | - | - | - |
| F124 | + | + | - | + | - | - | - | - | - | - | - | - | - |
| F135 | + | + | + | + | - | - | - | + | - | - | NA | NA | NA |
| F139 | - | - | NA | NA | - | - | NA | NA | - | NA | NA | + | + |
| F155 | - | + | + | + | + | + | - | - | - | - | NA | NA | NA |
| F292 | - | - | + | + | + | - | - | - | - | - | - | - | - |
| F326 | + | - | - | - | + | + | - | - | - | - | - | - | - |
| F335 | + | + | NA | NA | NA | NA | - | - | - | - | - | - | - |
| F487 | + | - | + | + | + | + | - | - | - | - | NA | NA | NA |
| F499 | + | + | - | - | - | - | - | - | - | - | - | - | - |
| F653 | + | + | - | - | - | - | - | - | - | - | - | - | - |

ACL: anticardiolipin. LAC: Lupus anticoagulant. B2GPI: Beta-2-glycoprotein I. NA: Not available

**Supplementary Table 4: Sequencing Metadata**. Each row is a sample. The sample name includes the individual subject, ANA status and B cell subset. Samples marked with asterisks only have a single sequencing replicate. Sequence copies refer to total reads in the library (pooled from both replicates). Clones = clone count from pooled replicates. Clones share the same VH, JH, CDR3 length and 85% amino acid sequence similarity in the CDR3. Additional filtering of sequences for clones is described in the Methods section. Mean CDR3 length is provided in nucleotides. Average V gene identity refers to the degree of nucleotide similarity to the nearest corresponding germline IGHV gene. A value of 1.0 would indicate a gene that is identical to the germline (unmutated). Somatic hypermutation (SHM) is (1-VH identity) x 100%. Top 20 refers to the percentage of sequence copies derived from the sum of the top 20 ranked clones. Total DNA input is the sum of the DNA amount in each replicate or in the only replicate for samples marked with an asterisk. Cell counts refers to the number of sorted cells.
